# Gene expression stasis and plasticity following migration into a foreign environment

**DOI:** 10.1101/121608

**Authors:** Brian K. Lohman, William E. Stutz, Daniel I. Bolnick

## Abstract

Selection against migrants is key to maintaining genetic differences between populations linked by dispersal. Yet, migrants are not just passively weeded out by selection. Migrants may mitigate fitness costs by proactively choosing among available habitats, or by phenotypic plasticity. We previously reported that a reciprocal transplant of lake and stream stickleback (*Gasterosteus aculeatus*) found little support for divergent selection. We revisit that experiment to test whether phenotypic plasticity in gene expression may have helped migrants adjust to unfamiliar habitats. We measured gene expression profiles in stickleback via TagSeq and tested whether migrants between lake and stream habitats exhibited a plastic response to their new environment that allowed them to converge on the expression profile of adapted natives. We report extensive gene expression differences between genetically divergent lake and stream stickleback, despite gene flow. But for many genes, expression was highly plastic. Fish transplanted into the adjoining habitat partially converged on the expression profile typical of their new habitat. This suggests that expression plasticity may soften the impact of migration. Nonetheless, lake and stream fish differed in survival rates and parasite infection rates in our study, implying that expression plasticity is not fast or extensive enough to fully homogenize fish performance.

## Introduction

What happens to an organism when it moves into a new habitat? Populations in disparate environments commonly exchange migrants. These migrant individuals are exposed to unfamiliar abiotic conditions and biotic communities to which their phenotypes may be poorly suited (Lenormand 2002; Kawecki and Ebert 2004; Hereford 2009). Migrants can be maladapted to their new habitat because they inherited alleles that were selectively favored in their native range but are untested by selection in their new habitat (Nosil, et al. 2005). Or, migrants’ traits may have been shaped, during ontogeny, by their native environment (Davis and Stamps 2004; Stamps and Davis 2006). Either way, migrants’ poor fit to their new habitat may frequently result in reduced survival, fecundity, or mating success (Hereford 2009). This selection against migrants is key to maintaining genetic differences between populations linked by dispersal (Lenormand 2002; Nosil, et al. 2005). Yet, migrants are not just passively weeded out by selection. Instead, migrants may evade selection in two ways. First, they can proactively choose among available habitats to avoid environments to which they are mismatched (Edelaar, et al. 2008; Edelaar and Bolnick 2012). Or, migrants may plastically alter one or more phenotypic traits to acclimate to a new habitat (Ghalambor, et al. 2007; López-Maury, et al. 2008; Davidson, et al. 2011).

Plasticity is most frequently measured as change in phenotype in response to an environmental change. Reciprocal transplants or common garden experiments have been successful at partitioning the relative contributions of heritability versus plasticity for a myriad of ecologically relevant traits (Conover and Present 1990; Pfennig 1992; Schlichting and Pigliucci 1998; West-Eberhard 2003). A meta-analysis of 258 studies on marine invertebrates strongly suggests that phenotypic plasticity in dispersal may evolve as a consequence of high habitat heterogeneity (Hollander 2008). A limitation of this literature, however, is a tendency to focus on readily-measured phenotypic traits (e.g., morphology and size), which may not be the most crucial traits for migrants’ fitness, and which may not be representative of plasticity for other more subtle yet important traits.

Gene expression profiling offers a much broader approach to assay the response of an individual to both abiotic and biotic stressors. Unlike phenomics, transcriptomics is not limited to a priori hypotheses regarding specific traits, and not only allows for trait but also pathway discovery. However, because of the substantial cost of transcriptomic analyses, there are few studies of transcriptome-wide plasticity in natural settings, and most of these have very limited biological replication (Todd, et al. 2016). Other studies have achieved higher replication (and thus power) by testing for plasticity of just a few candidate genes. For example, Stutz *et al*. (2015) showed that stickleback fish transplanted between lakes converged strongly to resemble the immune gene expression profile of natives of their new environment, indicating strong plasticity for a small set of seven genes (Stutz, et al. 2015). But, is this plasticity particular to immune genes, or is it representative of gene expression across the transcriptome?

Here, we describe how the stickleback transcriptome responds to an unfamiliar environment. We recently conducted a reciprocal transplant of threespine stickleback, moved between adjacent but contrasting lake and stream habitats. That study found little evidence of divergent selection: immigrants and residents grew and survived equally well, and had similar average parasite loads and diversity (Bolnick and Stutz in review, revised). Here, we present a study of the sticklebacks’ transcriptomic response to this transplantation. Specifically, we tested for i) baseline differences in gene expression between natives of each habitat, ii) differences in gene expression associated with being moved from one habitat to another, and iii) convergence in expression profiles between native and transplanted individuals.

The threespine stickleback fish (*Gasterosteaus aculeatus*) offers an opportunity to study plasticity of both phenotypes and gene expression. Across Vancouver Island, British Columbia, there are many replicate pairs of lake and stream stickleback (Thompson, et al. 1997; Hendry, et al. 2002). These parapatric lake and stream populations are typically genetically and phenotypically divergent (Roesti, et al. 2012; Feulner, et al. 2015; Weber, et al. 2017). These phenotypic differences persist to some degree in constant laboratory settings indicating there are heritable differences (Oke, et al. 2016)[for other common garden studies of phenotypic plasticity see (Kalbe and Kurtz 2006; Scharsack, et al. 2007; Berner, et al. 2011; Jiang, et al. 2016). Adjoining lake and stream environments differ in both abiotic and biotic conditions including flow regime, oxygen concentration, habitat structure, resource availability, prey composition, and parasite communities (Berner, et al. 2009; Kaeuffer, et al. 2012; Lenz, et al. 2013; Stuart, et al. in review, revised). The magnitude and direction of environmental differences between a lake and its outlet stream effectively predicts the direction of phenotypic differentiation between lake and stream resident stickleback (Stuart, et al. in review, revised). The implication, invoked by many studies of lake and stream stickleback (summarized in (Weber, et al. 2017)), is that environmental differences drive divergent selection on lake and stream stickleback.

To test for this inferred selection, multiple studies have transplanted lake and stream stickleback into their native and neighboring habitats, measuring whether residents systematically outperform immigrants in a variety of measures (survival, growth, infection)(Hendry, et al. 2002; Bolnick 2004; Scharsack, et al. 2007; Hanson, et al. 2016; Moser, et al. 2016; Kaufmann, et al. 2017; Bolnick and Stutz in review, revised). However, these experiments yielded surprisingly inconsistent evidence for divergent selection (summarized in extended data of (Bolnick and Stutz in review, revised)). Why is divergent selection rarely observed (but see (Hendry, et al. 2002; Kaufmann, et al. 2017), despite evidence of phenotypic divergence? Several recent papers discuss the possibility of habitat choice helping to maintain lake-stream differences (Bolnick, et al. 2009; Berner and Thibert-Plante 2015; Jiang, et al. 2015; Weber, et al. 2017). Another possibility is that plasticity mitigates selection against migrants. Here, we use a reciprocal transplant experiment that found negligible support for divergent selection, to also test for plasticity. We measured both physical traits (e.g. change in mass over a given period or the value of an ecologically relevant trait) and gene expression profiles via RNAseq (Lohman, et al. 2016) for a large number of transplanted individuals. Using this data, we tested whether migrants’ gene expression shifts to more closely resemble expression by the native population in their new habitat, suggesting a role for expression-mediated phenotypic plasticity by migrants.

There is ample evidence for phenotypic plasticity in ecologically relevant traits in stickleback. For example, previous experiments reared stickleback from different habitats in a common garden setting (lab aquaria), and fed them alternative diets to test for plasticity in feeding morphology (Day and McPhail 1996; Svanbäck and Schluter 2012). These studies measured body shape, gill raker, and gape traits that are both readily measured and clearly relevant to foraging. Life history traits also show plasticity in stickleback, including breeding size, clutch size, egg size, and relative clutch mass (Baker and Foster 2002). Finally, prior studies have examined plasticity in gene expression (Wang, et al. 2014; Leder, et al. 2015; Robertson, et al. 2015; Gibbons, et al. 2017). One such study focused on expression of two candidate genes for osmoregulation and salinity tolerance (McCairns and Bernatchez 2010). A larger, whole transcriptome approach suggested that the invasion of freshwater and thermal tolerance drove the evolution of gene expression plasticity (Morris, et al. 2014). However, while these studies of gene expression plasticity have sought to answer how the transcriptome may respond to a novel environment, they have been carried out in the lab and do not account for the diverse stressors of the wild. We therefore tested whether migrants between lake and stream habitats indeed exhibit a strong plastic response to their new environment that allows them to converge on the gene expression profile of the adapted natives.

## Methods

### Sample acquisition

As detailed in (Bolnick and Stutz in review, revised), stickleback from Roberts Lake and stream (Vancouver Island, British Columbia, Canada) were trapped, weighed, measured for length, individually marked with unique spine clips, then placed in cylindrical wire cages. Lake cages and stream cages both received a total of 60 lake fish and 60 stream fish. Each cage was approximately 1.6m in diameter, placed in 1m deep water and sealed to the substrate to prevent escape. Cages were made of wire mesh that allowed free flow of water and movement of prey items. In the lake, cages were situated along the shoreline roughly 150m from the outlet stream. In the stream, cages were placed 150m downstream from the lake. In an effort to reduce the influence of gene flow on stream genotypes, stream fish for the experiment were gathered from 1.5 km upstream of the lake outlet. Each enclosure contained 3 fish, half the cages receiving a 1:2 ratio of lake:stream fish, the other half of the cages receiving a 2:1 ratio. Within each cage, the three fish were uniquely marked with dorsal spine clips to facilitate identification. After 8 weeks, we recaptured the caged stickleback. As a control for the effect of caging we also collected wild uncaged fish from both lake and stream at the conclusion of the experiment, from habitat immediately adjoining the cages. Hereafter, here we refer to uncaged fish as the ‘wild’ group, all fish recovered from cages are ‘transplanted’. Within the transplanted fish we distinguish between ‘natives’ (same origin and destination habitats) and ‘immigrants’ (different origin and destination).

We euthanized the collected fish with an overdose of MS-222. We weighed each fish, measured its length, and dissected the fish to remove head kidneys (‘pronephros’) which we stored in RNAlater (Ambion) for subsequent RNA extraction and expression analysis. After dissection, specimens were preserved in ethanol for later dissection for to enumerate parasites. We sequenced MHC IIb exon 2 from all fish (using DNA from pre-release spine clips), as described in (Stutz and Bolnick 2014; Bolnick and Stutz in review, revised).

### RNAseq library preparation, sequencing, and bioinformatics

Following total RNA extraction (Ambion AM1830) we built RNAseq libraries according to (Lohman, et al. 2016). TagSeq libraries were sequenced on the HiSeq 2500 with 1x100 V4 chemistry at the Genome Sequencing and Analysis Facility at the University of Texas at Austin, generating an average of ∼5 M raw reads per sample.

Raw reads were processed according to the iRNAseq pipeline (Meyer, et al. 2011; Dixon, et al. 2015; Lohman, et al. 2016), producing a total of 19,556 genes. Due to a machine error during the HiSeq run, BaseSpace was unable to convert cycle 35 to a base call, and thus base 35 is N in every read. We adjusted for this by adding the –n option to all calls to fastx_clipper in the iRNAseq pipeline. Mapping with Bowtie2 should not be influenced by this error (∼53.3% alignment rate, post quality filtering, adaptor trimming, and poly-A removal). GO enrichment was conducted according to (Wright, et al. 2015) using transcriptome annotation built with the UniProtKB database and following previously described procedures (Dixon, et al. 2015; Lohman, et al. In review).

### Statistical Analysis

We analyzed gene expression using a series of linear models in DESeq2 (Love, et al. 2014), limma (Ritchie, et al. 2015), and base R (R Development Core Team 2007). We sought to estimate three effects:

### 1. What are the differences between wild fish from Roberts Lake and Stream?

We tested for differences in gene expression between wild (uncaged) fish from Roberts Lake versus Roberts Stream by modeling gene count as a function of origin (lake or stream). We tested for GO enrichment within the main effect of origin with a Mann-Whitney U via GO_MWU (Dixon, et al. 2015). We used weighted gene coexpression network analysis (WGCNA; (Langfelder and Horvath 2008) to estimate correlations between suites of coexpressed genes and traits, including morphology, parasite burdens, and genotypes (e.g. MHC allelic diversity). WGCNA is an unbiased, data-driven method to cluster groups of genes with similar expression patterns. We removed batch effects and normalized counts using limma (Ritchie, et al. 2015) before starting WGCNA. We followed the tutorial of (Langfelder and Horvath 2008), and constructed a signed network with a soft thresholding power of 6 and a minimum module size of 30 genes. We used dynamic tree cut and merged modules with greater than 80% similarity, producing a total of 11 modules. We plot the FDR corrected Pearson correlation coefficient between module eigengenes and trait values.

### 2. What is the effect of being transplanted into a novel environment?

We tested for changes in gene expression of transplanted (caged) fish as function of origin habitat, destination habitat, and the interaction between origin and destination. Using our estimated gene network, we calculated the FDR corrected Pearson correlation coefficient between module eigengenes and traits unique to the transplant design (treatment, origin, destination, delta mass, and delta length). A main effect of origin indicates stable gene expression differences between native lake versus native stream fish. These expression differences can be stable because they are heritable, or because they are environmentally-induced only early in ontogeny but remain canalize in adults, which we used for this experiment. A main effect of destination indicates genes that respond plastically to recently experienced environments. An interaction between origin and destination would indicate ecotype differences in how they respond to a given environment. Such interactions could entail G*E effects on expression, but we point out they could also arise from ecotype differences in the extent of canalization of early plasticity.

### 3. How well do immigrants converge on the expression profile of natives?

We conducted a PCA of expression of all genes in all fish, then selected only transplanted fish and used the leading # PC axes for subsequent linear discriminant analysis. The original expression matrix has too much collinearity for LDA. Dropping higher-order PCA axes reduces this collinearity, enabling LDA. This approach is sometimes called DAPC (Jombart, et al. 2010; Kenkel and Matz 2016). We plotted these results in LDA space, adding vectors connecting each ecotype’s expression at home to the same ecotype’s expression in the foreign habitat. These vectors represent the magnitude and direction of expression plasticity along DAPC axes. Convergence in expression would result in an angle of 180 degrees between the vectors for lake and stream ecotypes. Moreover we compared the lengths of these vectors to evaluate whether lake and stream ecotypes are equally plastic.

Lastly, if plasticity effectively recreates lake-stream differences then we would expect that genes that are more highly expressed in lake natives would also be more highly expressed in fish transplanted into the lake. This can be tested by measuring the correlation, across genes, between the origin effect sizes and destination effect sizes estimated in analysis (2) above. Adaptive plasticity to converge on a native expression profile should result in a positive correlation.

## Results

### What are the differences between wild fish from Roberts Lake and Stream?

Our linear model revealed that 647 genes were differentially expressed between wild Roberts Lake and Roberts stream stickleback (Wald, p < 0.1 after 10% FDR, or 306 when p < 0.05). GO analysis showed that these genes are enriched both for a variety of categories including (but not limited to) genes regulating macrophage differentiation (biological processes, Mann-Whitney U, p < 0.05 after 10% FDR correction, Figure 1), and genes involved in the MHC Class II protein complex (cellular components, Mann-Whitney U, p < 0.1 after 10% FDR). Both of these GO groups have known functions in parasite defense, and have been previously implicated in the response to selection and parasite prevalence in stickleback (Lohman, et al. In review; Bolnick and Stutz in review, revised). Previous studies revealed that Roberts Lake and Stream stickleback populations harbor significantly different parasite communities (Bolnick and Stutz in review, revised), with corresponding differences in MHC Class II allele frequencies (Stutz and Bolnick In review).

**Figure 1.**
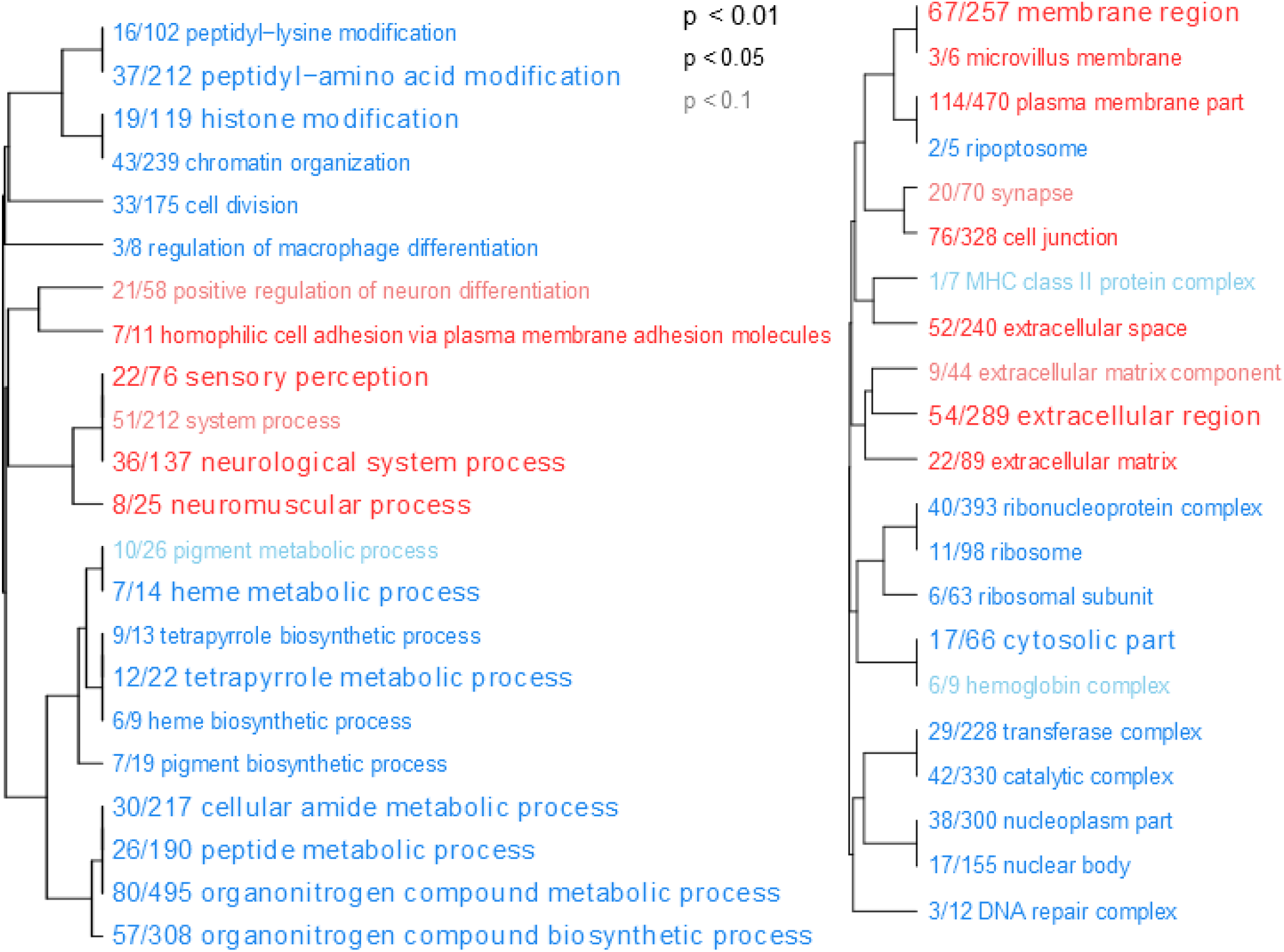
GO analysis results on Lake vs Stream wild fish. Blue terms are underexpressed while red terms are overexpressed relative to the lake baseline. P-values are Mann-Whitney U. Dendrograms indicate similarity of GO groups. Left group is from the Biological Processes cluster while the right group is from Cellular Components.

In addition to gene-by-gene linear modeling we also tested for correlations between modules of coexpressed genes and various traits, including morphology, infection by parasites, and MHC Class IIb genotype (Figure 2). We found morphology to be correlated with many different modules, each with modest correlation but highly significant p-values. It is noteworthy that all modules except the ‘turquoise’ module have a negative correlation with morphology (co-expressed gene modules are given arbitrary color names). There are correlations between MHC allele number and several modules, including greenyellow, blue, magenta, and pink. Interestingly, MHC allele number and two measures of parasite diversity have equal strength but opposite sign in their correlation to the greenyellow module (Figure 3A). This is consistent with previous experimental and theoretical data that animals with more diverse MHC genotypes should have fewer parasites (Wegner, et al. 2003). Finally, we also considered linear discriminant axes of MHC II genotypes from a prior analysis of these same fish. We find that LDA1 and LDA3 of MHC II are correlated with turquoise and purple modules. The turquoise expression module is also correlated with fish origin (r = 0.36, p << 0.001) so these correlations are likely a result of differences in MHC genotype between the two ecotypes. Infection by several functional groups of parasites are significantly correlated with particular modules. For instance, the purple module is correlated with infection by nematodes (r = -0.22, p < 0.04, Figure 3B) and genes in the red module are correlated to infection by any species of *Proteocephalus* (Figure 3C, r = 0.28, p < 0.01).

**Figure 2.**
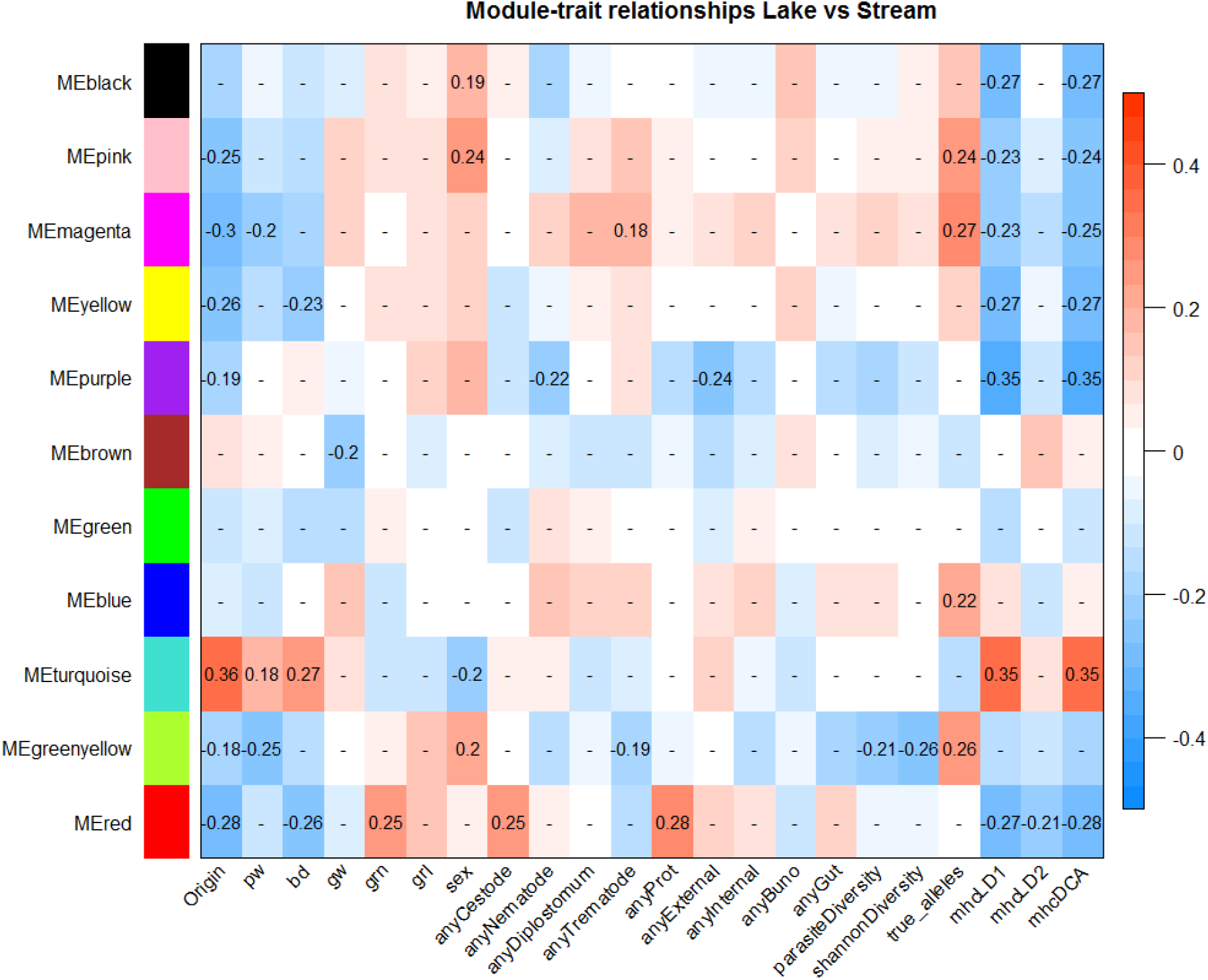
WGCNA reveals correlations between modules of coexpressed genes and traits in wild lake and stream fish. Cell values are Pearson correlation coefficients. Only correlations with p-values less than 0.1 are presented. Modules shown are the same as Figure 5.

**Figure 3.**
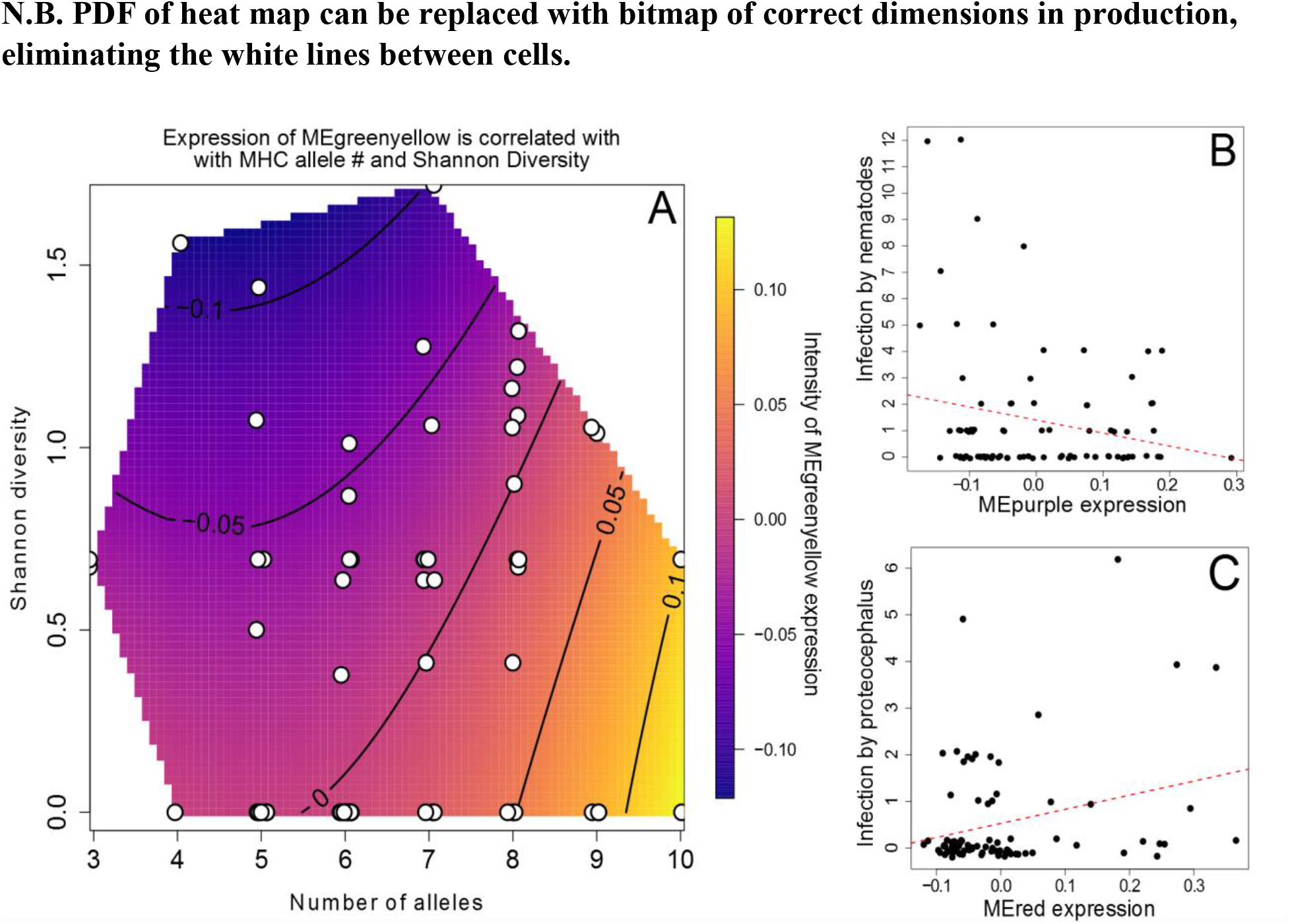
Module trait correlations from WGCNA. A) Heat map of MEgreenyellow expression shows negative correlation with Shannon diversity of parasite infection and positive correlation with MHC allele number. B) Increased expression of MEpurple is correlated with decreased infection by nematodes. C) Increased expression of MEred is correlated with increased infection by proteocephallus.

### What is the effect if being transplanted into a novel environment?

Focusing next on transplanted (caged) stickleback, we observed significant effects of both origin and destination for many genes (507 and 111, respectively when q < 0.05, see supplementary material for full list). Here, the effect of origin represents genotype effects that persisted after transplantation (because the effect of transplantation is averaged). Approximately 94% of the genes with significant (q < 0.1) origin effect in transplanted fish were also significantly different between wild fish ecotypes. This overlap of origin effects in caged and wild fish suggests that stickleback exhibit realistic lake-stream expression differences when placed in lake or stream cages.

Destination effects represent plasticity that was independent of genotype (genotype effects are averaged in our model). Most notably, this list of genes includes *hsp90* (lower in fish transplanted into the stream, Wald, p << 0.001 after 10%FDR), a stress response protein which has been studied in many different animals (Rutherford and Lindquist 1998; Queitsch, et al. 2002). In addition, *stat1* (lower in fish transplanted into the stream, Wald, p < 0.09 after 10% FDR), was also significantly different between fish transplanted in alternate environments. This transcription factor has a rich history of study for its critical role in multiple signaling cascades throughout the immune system (Murphy 2011).

We found only 10 genes whose expression depended on the interaction of origin and destination (Wald, p < 0.1 after 10% FDR, or 4 when p < 0.05, Figure 4, supplementary Table 1, Supplementary Figure 1). Such interactions can loosely be interpreted as genotype by environment interactions (e.g., genetic differences in plasticity), with the caveat that we are studying wild-caught fish. Of these 10 genes, two candidates are possibly involved in defense against parasites: *cyp24a1*, a cytochrome p450 variant (Annalora, et al. 2010) and *dhx58*, an antiviral gene about which little is known (Leavy 2012). In both cases lake natives have higher expression in the lake than do stream fish, but decreased expression when moved into the stream. Stream fish have higher expression in their native habitat, but only higher than foreign lake fish for *dhx58*. Furthermore, it is noteworthy that for all 10 interaction genes, the genes are more highly expressed in lake than in stream fish (all in lake cages). And, for all 10 genes the lake natives decrease expression when moved into the stream (Figure 4).

**Figure 4.**
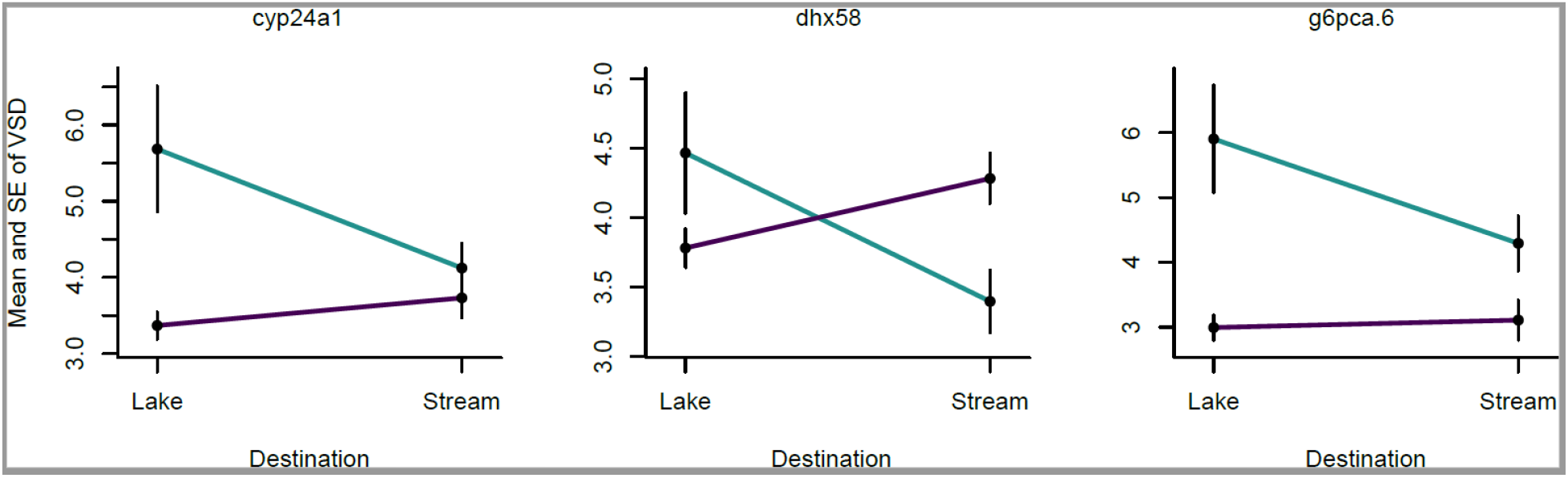
Reaction norm plots of genes significant for interaction between origin and destination. X-axis is destination, teal line is lake and magenta line is stream origin/genotype. Vertical line indicate standard error.

We used DESeq2 to estimate caging effects by comparing wild fish to natives within each environment. We found a moderate number of differentially expressed genes. 35 genes were differentially expressed between wild lake fish and caged lake natives. Somewhat more genes (79) were differentially expressed between wild stream fish and stream natives (all Wald, p < 0.1 after 10% FDR, or 19 and 52 when p < 0.05, respectively. See supplementary material for full list). There are very few notable differences due to caging in lake genotypes. Lake natives have higher expression of *cyp24a1* than wild lake fish (log2 fold change = 3.8, p = 0.065 after 10% FDR correction). Lake transplants also have higher expression of *ebf4*, an early B-cell factor (log2 fold change = 4.3, p = 0.049 after 10% FDR correction) than wild lake fish. In contrast, when we make the same contrast but in stream genotypes, almost all differentially expressed genes (76 out of 79 passing p < 0.1 after 10% FDR correction) exhibit a pattern of lower expression in transplants than in wild fish (see supplementary material for full list of genes and statistics). Stream transplants have lower expression of immune genes with known function including the complement system (complement 3, 8, and 9), a leukocyte derived chemotaxin (*lect2l*), and three fibrinogen genes (alpha, beta, and gamma). In addition, two coagulation factor genes are lower in natives (factor 13 and 7i) then wild stream fish. The cage effect for stream fish is partially confounded, however, with genotype. The stream transplants were from 1.5 km downstream of the cage site, whereas the wild fish were collected among the cages, 100m downstream from the lake. So, differences between wild stream and transplanted stream fish may be genetic rather than exclusively a plastic response to caging. There were almost no genes (only 2) that showed significant effects of caging in both the lake and in the stream, indicating that there is no generic transcriptomic response to caging (Supplementary Figure 2).

Our coexpression analysis of transplanted fish revealed significant correlations between traits unique to this subset of fish and modules of gene expression. For example, treatment (transplanted into foreign or native environment) and origin are both correlated with the turquoise module. In contrast, destination is only weakly correlated to the pink and magenta modules. The red module has a negative correlation to origin and a positive correlation to change in length over the course of the experiment. Change in mass is correlated to both the magenta and purple modules. Interestingly, there is no overlap between change in mass and change in length. This difference suggests a change in condition within individuals (Figure 5).

**Figure 5.**
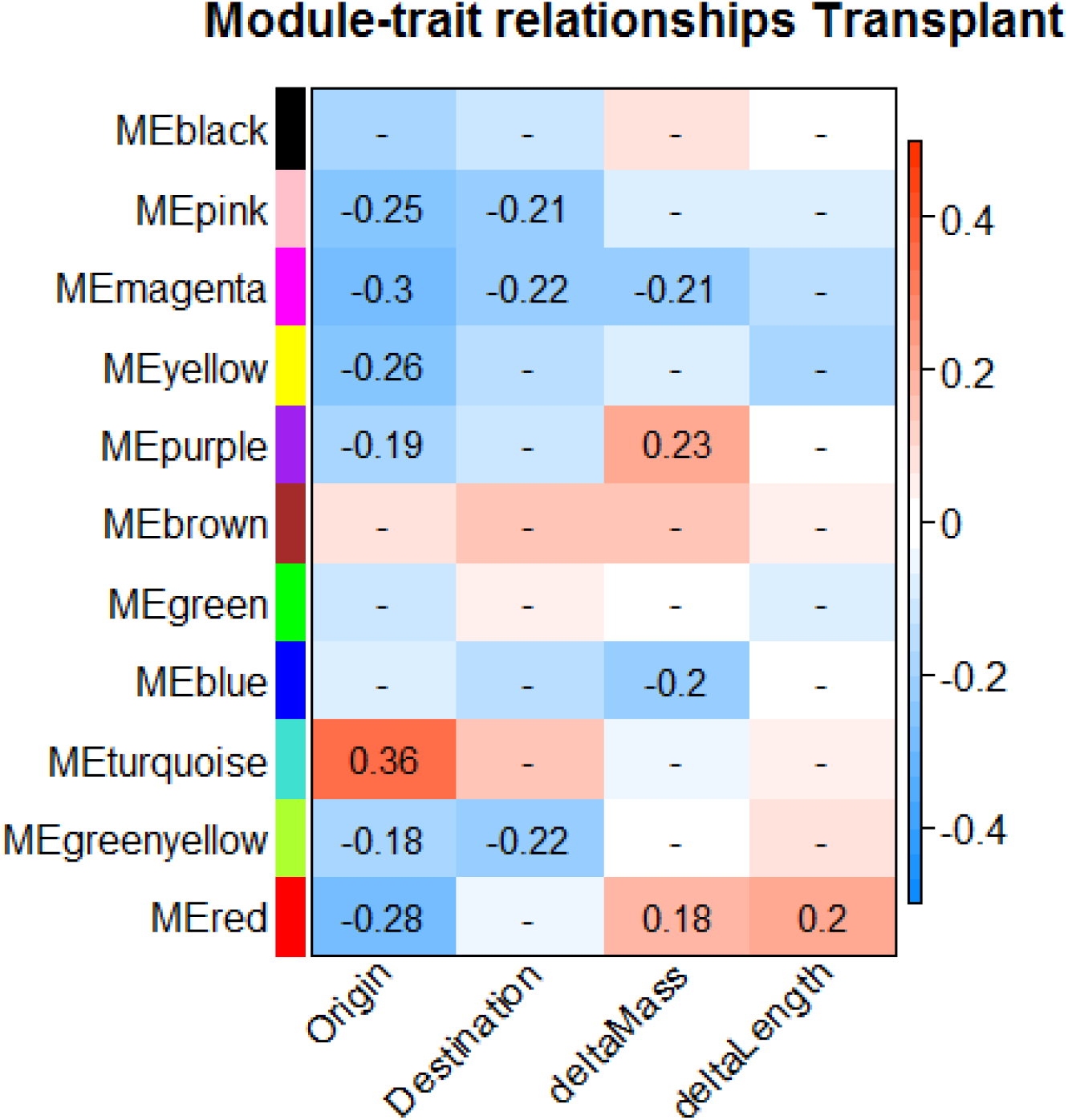
WGCNA reveals correlations between suites of coexpressed genes and traits in transplanted fish. Cell values are Pearson correlation coefficients. Only correlations with p-values less than 0.1 are presented. Modules shown are the same as Figure 2.

### How well do immigrants converge on the expression profile of wild controls?

We tested for convergence between natives and immigrants in the entire expression profile. Within a bivariate discriminant function space, we found that LDA1 separates fish by origin (lake versus stream, explains 86% of variance). LDA2 separated fish based on their transplant destination (explains 10% of variance). LDA3 roughly separates native/non-native status (, explains 3.5% of variance, Figure 6 and supplementary Figure 3). We plotted a vector from the mean of each resident ecotype at home, to the mean expression of the same ecotype when moved into a new environment. The vector showing the expression change of lake fish is almost in exactly the opposite direction from the expression change of stream fish (∼180 degrees, visually). In each case, fish moved into a new habitat converged on the expression profile of their new neighbors along LD2 (but not along LD1 or LD3). Lake fish moved into the stream actually overshot the stream expression profile, resulting in a much larger reaction norm vector than stream fish moved into the lake (LD2 ∼origin + destination + origin:destination) and found a significant effect of the interaction (p << 0.001). Because of this overshooting, both the lake-to-stream migrants and stream-to-lake migrants were significantly different (for LD2) from the resident ‘target’. We conclude that immigrant stickleback partially converge on native expression profiles after emigration to a new habitat, and that lake fish exhibit stronger plasticity. The latter finding matches the greater plasticity of lake fish in our gene by gene analysis with DESeq2 (above; Figure 7).

**Figure 6.**
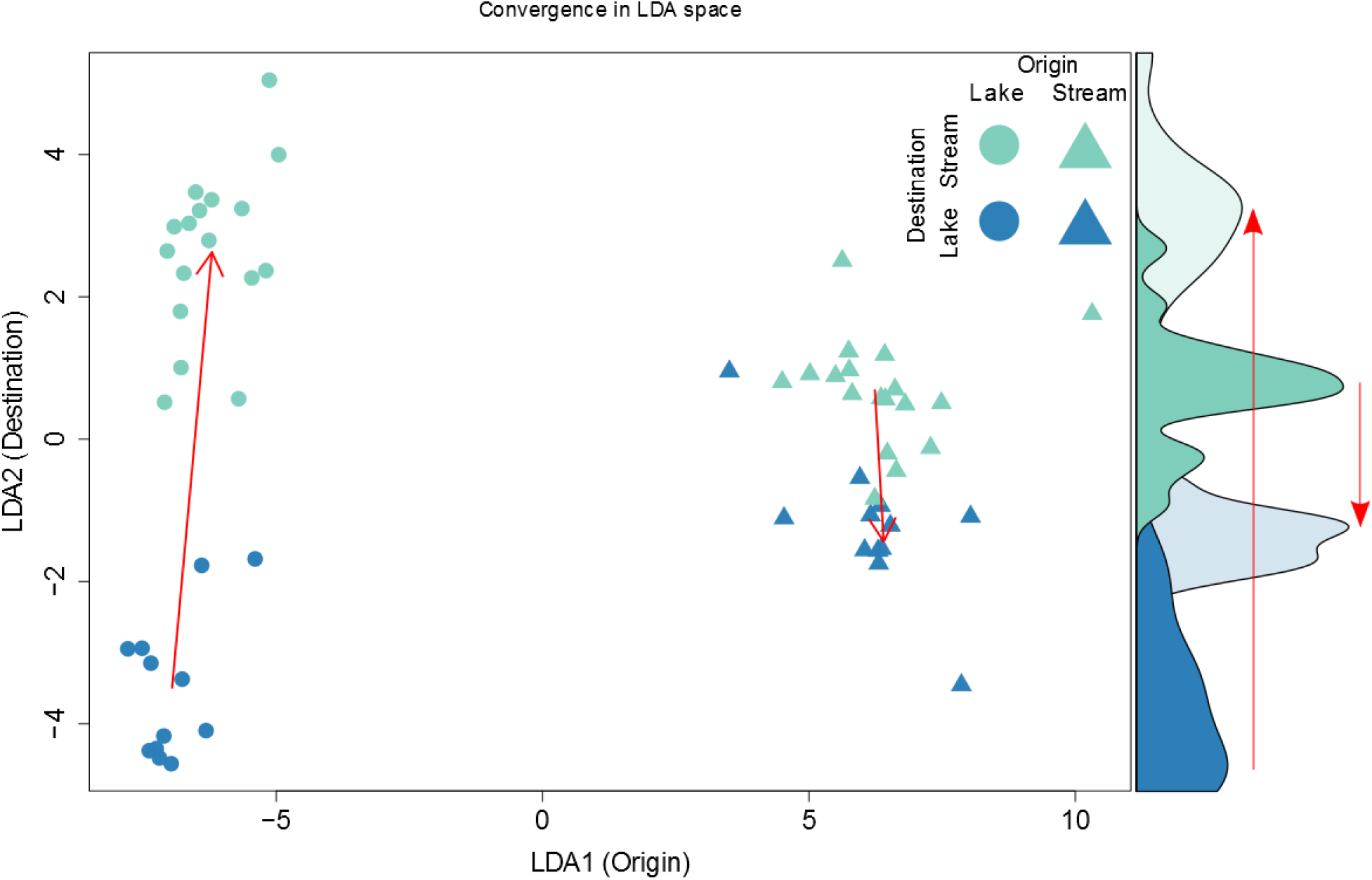
Convergence of immigrant expression profiles toward native expression profiles in transplanted fish. Red arrows are drawn between the means of each distribution. Fish originating from the lake move farther along LD2 than stream fish (two factor ANOVA, p << 0.001).

**Figure 7.**
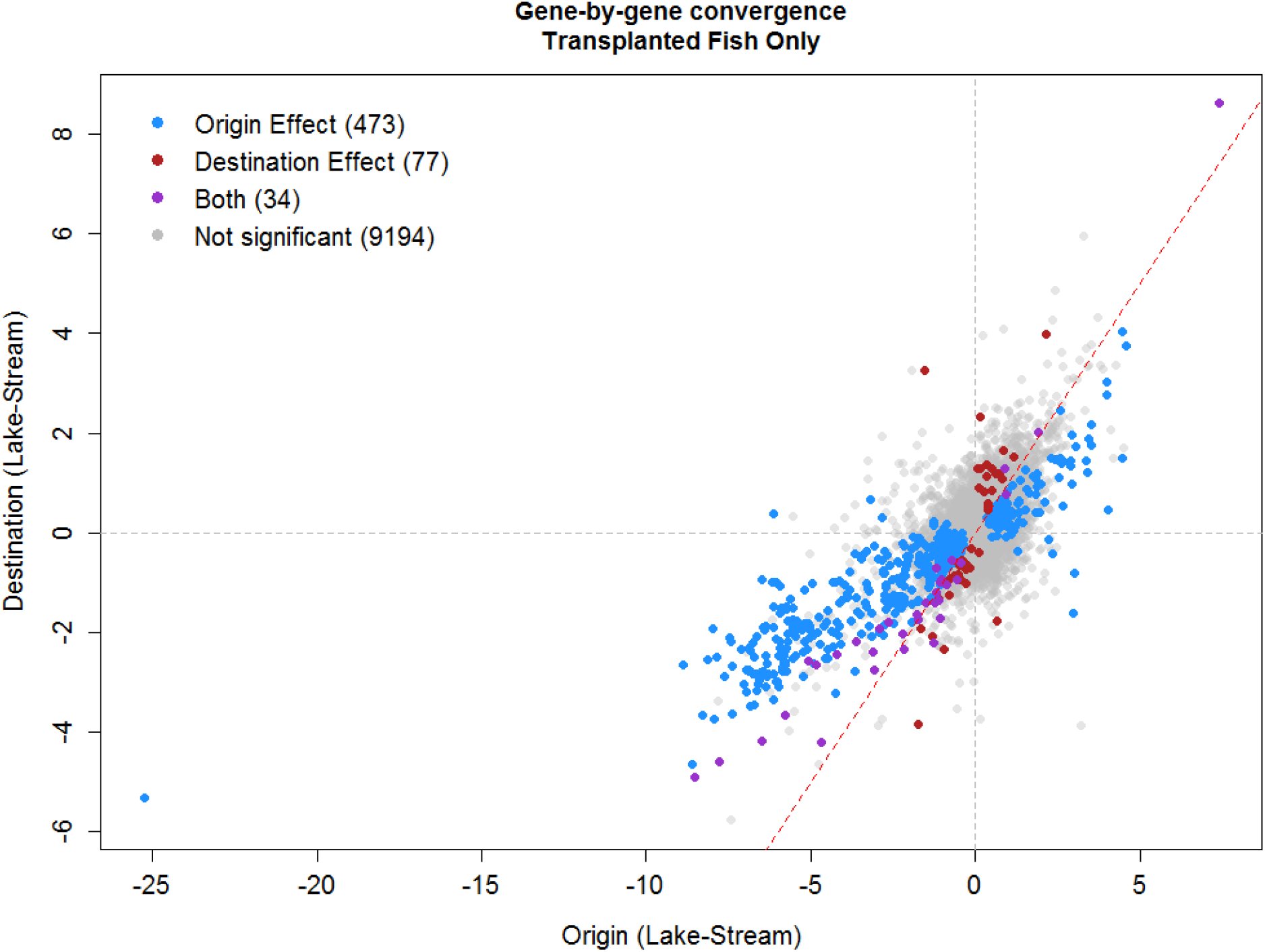
Gene-by-gene convergence among transplanted fish. We included only transplanted fish in a linear model in DESeq2: with expression of each focal gene as a function of origin + destination + origin:destination. X and Y axis are Log2 fold changes between lake and stream fish by origin, and destination, respectively. Points are colored when q-value < 0.05, and color coded based on which effect(s) were significant. The red dashed line is 1:1, helping to visualize that the main trend has a slope less than 1, indicating that plasticity (destination) effects are weaker than origin effects.

We also considered convergence at the individual gene level. Using the DESeq2 linear model estimates, we found that destination effects were positively correlated with origin effects (r = 0.67, Figure 7). That is, transcripts that were more abundant in lake natives were also more abundant in fish placed in lake cages, and vice versa for stream-biased transcripts (Figure 7). This implies that for many genes, expression differences between the native populations are recapitulated by plastic responses to animals’ recent environment. The observed destination origin relationship has a slope less than 1 (0.42, p << 0.001) indicating that the plasticity is not, however, complete, which fits with the fact that the major LDA axis still separates lake versus stream natives and explains more variation than the second LDA axis that measured plasticity.

## Discussion

Organisms’ adaptation to their native habitats means that migrants will often be maladapted to novel environments. One way that migrants may be able to ameliorate stressors of new habitats is by modulating gene expression. Prior studies have used reciprocal transplants to uncover plasticity in select candidate genes, but this approach could miss a myriad of responses to the environment. To look for static and plastic responses in gene expression associated with emigration, we tested for differences in gene expression among stickleback reciprocally transplanted between two adjoining habitats containing genetically divergent populations. We found expression differences between these populations, and changes in response to emigration, at the level of individual genes, gene coexpression, and the whole transcriptome.

### There are constitutive differences in gene expression between lake and stream stickleback

Although Roberts Lake and stream are adjoining habitats that permit easy movement of stickleback between sites, the resident stickleback populations are genetically distinct. Fish from this lake and stream differ in a range of morphological and parasitological traits (Berner, et al. 2009; Oke, et al. 2016; Weber, et al. 2017; Stutz and Bolnick In review; Bolnick and Stutz in review, revised), as is true for many such lake-stream pairs (Stuart, et al. in review, revised). Given these genetic and phenotypic differences, we expected to find differences in gene expression between these populations. Roughly 7% of the 9748 genes in our transcriptome dataset exhibited between-population differences in relative abundance.

Some of these differences fit well within the existing literature of lake-stream divergence. For example, our GO enrichment results suggest that macrophage differentiation is different between lake and stream fish. Macrophages contribute to initiation of immune defenses against a variety of parasites including but not limited to the tapeworm *Schistocephalus solidus* (Kurtz, et al. 2006), whose infectious procercoids are deposited by loons and mergansers (which prefer lakes over streams) and carried by zooplankton (which are more abundant in lakes than streams). MHC class II is another parasite defense related GO category which is different between lake and stream. Prior work in the Roberts lake stickleback has revealed that MHC II allele frequencies differ between this particular lake and stream (Stutz and Bolnick 2014), as well as many other lake-stream pairs (Kurtz, et al. 2006; Wegner, et al. 2006; Eizaguirre, et al. 2010). Furthermore, individuals who carry local MHC alleles are more heavily infected with parasites than individuals carrying foreign MHC alleles (Bolnick and Stutz in review, revised). Our WGCNA results suggest that MHC allele diversity and parasite diversity are negatively correlated with each other and jointly associated with a set of co-expressed genes. Specifically, the greenyellow module has a negative correlation with parasite diversity and a positive correlation with the number of MHC alleles (Figure 2, 3A). While this result stands out as support for a large body of theory (Eizaguirre and Lenz 2010; Spurgin and Richardson 2010) and agrees with prior empirical evidence (Wegner, et al. 2003; Piertney, et al. 2009), we would have expected greater correlations between modules and parasite infection. However, this lack of correlation is likely due to sparse and overdispersed parasite infections, which make correlations difficult to estimate well.

### Transplantation into alternate habitats reveals static and plastic gene expression

For our experimentally transplanted fish, individuals’ origin accounted for more expression variation (507 genes) than did destination (111 genes, Figure 7). The main effect of origin represents persistent between-population differences no matter which habitat the fish were caged in. Thus, we interpret the main effect of origin as a probable signal of static genetic differences in expression, insensitive to the environment. However, we also found significant destination effects for a subset of genes, indicating appreciable plasticity in gene expression in response to sticklebacks’ recent (cage) environment. That is, expression of certain genes was higher in lake-caged fish than stream-caged fish, regardless of their origin. This plasticity is consistent with prior evidence that morphological plasticity contributes to phenotypic differences between the Roberts Lake and Stream stickleback (Oke, et al. 2016).

Notably, there was a positive correlation between origin effect and destination effect (r = 0.67). We therefore infer that the heritable lake to stream differences were at least partly recapitulated by plasticity. Genes more highly expressed in lake (stream) natives were also up-regulated in all fish placed in lake (stream) cages (Figure 7). If expression was exclusively plastic (on the time-scale of our experiment), we would expect to see no origin effect at all, which is not the case. So, this correlation between origin and destination effects suggests that heritable and plastic differences jointly contribute, in the same direction, to between-ecotype differences in expression. The fact that the origin-destination effect correlation has a slope less than 1 confirms the statement, above, that heritable (origin) effects were somewhat stronger than the environmental (destination) effects. Moreover, the paucity of genes in the top-left and bottom-right quadrants of Figure 7 suggests that remarkably few genes exhibited plastic responses that opposed the heritable lake-stream differences.

Very few genes (10) were significant for the interaction of origin and destination. This is consistent with prior observations that there are no interactions effects between origin and destination for parasite load, survival, growth, or condition in this experiment (Bolnick and Stutz in review, revised). The few interactions that do exist follow two distinct patterns. First, some genes were down-regulated after individuals were placed in a foreign habitat. Second, other genes were more highly expressed by lake fish, but also showed stronger plastic down-regulation in lake fish placed in the foreign stream habitat. The absence of genes which were more highly expressed by stream fish, regardless of habitat, is notable. Some of the interaction genes we do detect may be involved in ROS production and antiviral response, both of which may be potentially important to fitness. For example, ROS production was recently shown to be a heritable response to infection by *S. solidus* (Weber, et al. in review, revised).

We observed no cases where expression was higher in foreign habitats. While this could be a product of the low number of interaction genes, this pattern is surprising and worth considering in future work. Intuitively we would have expected transplanted fish in either direction to up-regulate stress genes, but this apparently did not occur. Perhaps the absence of interaction effects on genes here is because plasticity reinforced between-ecotype differences. The paucity of interaction effects may also be a consequence of our analytical technique: log-transformation of expression levels converts multiplicative (interaction) effects into additive effects, which can reduce power or even completely obscure our ability to detect significant interactions between origin and destination effects. Nevertheless, other reciprocal transplant studies using large scale RNAseq have found more interaction genes and this seems to be relatively common (Lovell, et al. 2016; Reid, et al. 2016).

Our WGCNA analysis revealed only weak correlations between origin and phenotypes unique to transplanted fish. For example, change in mass and length. However, it is interesting to note that changes in mass and length are most correlated with different modules. This may suggest a change in condition (loss of mass but increase in length due to growth but poor foraging efficiency, Figure 5).

### On the whole transcriptome level, lake fish are more plastic than stream fish

At the whole-transcriptome level, we again observe substantial and persistent differences between the expression profiles of lake and stream fish, captured by LD axis 1. However, along LD2 we observe substantial plastic convergence of immigrant fish towards the expression profile of their new population (Figure 6). Interestingly, we also observed convergence in parasite community composition in this same experiment. Lake and stream natives carried distinct parasite communities, and individuals transplanted to the neighboring habitat exhibited an intermediate parasite community (Bolnick and Stutz in review, revised).

Our analysis suggests that fish from the lake exhibit a more plastic response to being transplanted into the stream, compared to stream fishes’ more limited plasticity when placed in the lake. This transcriptome-wide analysis is consistent with our single-gene analyses which also found that lake natives tended to show greater plasticity in response to transplantation. Assuming fish caged in their native habitat adopt a locally optimal expression profile, we infer that lake sticklebacks’ strong plastic response is actually excessive, overshooting the stream profile along LD2. In contrast, stream fish placed in lake cages fall short of the optimum expression in the lake (Figure 6). We therefore conclude that transcriptomic plasticity is incomplete (LD1 remains intact and explains the most variance), and differs between lake and stream ecotypes. This result implies that sticklebacks’ transcriptional reaction norms may be evolving as they adapt to different habitats. However, because we used wild-caught rather than lab-raised fish for this experiment we cannot rule out effects of early rearing environment, and hence cannot definitively ascribe a genetic cause to the different reaction norms of lake and stream fish.

Our results lend additional support to an emerging insight, that transcriptomic plasticity plays a substantial role in migrants’ adaptation to novel environments. This has been very extensively explored in experimental settings in the laboratory, where organisms may be exposed to alternative environmental conditions (often a single variable such as salinity, temperature, or a toxin). Many studies find plastic responses in candidate genes, or a subset of the transcriptome, in response to such experimental treatments (Whitehead, et al. 2011; Morris, et al. 2014; Reid, et al. 2016; Velotta, et al. 2016). Often these plastic responses are genotype-dependent, with one population exhibiting a stronger response than another (e.g., PCB-tolerant killifish are less plastic than PCB-susceptible populations (Reid, et al. 2016)). Fewer studies have examined transcriptomic plasticity of migrants in natural settings. Kenkel and Matz (2017) subjected corals to a reciprocal transplant experiment across a temperature gradient, and also found transcriptomic convergence of migrants towards residents, as we do. They also found that one genotype was more transcriptionally plastic than the other, as we do.

A large body of existing empirical and theoretical studies suggest that increased plasticity should evolve in more temporally or spatially heterogeneous habitats (Dudley and Schmitt 1996; Van Buskirk 2002; Auld and Relyea 2011; Baythavong 2011; Davidson, et al. 2011; Murren, et al. 2015). Our result is thus somewhat puzzling, in that we observe greater transcriptomic plasticity in lake fish, which inhabit the more temporally stable habitat. While stream habitats are generally very diverse (flow regime, overhead foliage, substrate, spatial distribution of prey), lake habitats generally have large and smooth transitions between any variation in environmental variables (and in most cases very little variation (Ahmed, et al. in press; Stuart, et al. in review, revised). However, lakes may be less predictable in other ways. For instance, lake stickleback consistently harbor more diverse parasite communities (Stutz and Bolnick In review; Bolnick and Stutz in review, revised), and so may have evolved greater immunological plasticity to handle an unpredictable suite of pathogens and helminthes.

In conclusion, we see extensive gene expression differences between genetically divergent stickleback populations inhabiting adjoining habitats but connected by gene flow (Weber, et al. 2017). But, for many genes, transcript abundance is highly plastic. Fish that disperse into the adjoining foreign habitat will partially converge on the gene expression profile typical of their new habitat. This suggests that expression plasticity can soften the impact of immigration into an unfamiliar habitat. Nonetheless, lake and stream fish differed in survival rates and parasite infection rates in our study, implying that this expression plasticity is not fast or extensive enough to fully homogenize the lake and stream fish performance.

## Acknowledgements

Collection and animal handling were approved by the University of Texas Institutional Animal Use and Care Committee (Protocol # 07-032201), and a Scientific Fish Collection Permit from the Ministry of the Environment of British Columbia (NA07-32612). We wish to thank Chad Brock and Kelsey Jiang for their assistance with collecting stickleback head kidneys during the reciprocal transplant experiment. The staff of the Genome Sequencing and Analysis Facility at the University of Texas at Austin provided technical support. This work was supported by the Howard Hughes Medical Institute (DIB) and the David and Lucille Packard Foundation (DIB)

## Data Accessibility

Meta data, parasite data, code for processing raw reads, code for statistical analysis and plotting is located in DRYAD entry [XXXX]. Raw reads are available for download via ‘wget [XXXX]’ from the terminal. The iRNAseq pipeline is available here: <u>https://github.com/z0on/tag-based_RNAseq</u>. The GO_MWU software is available here: <u>https://github.com/z0on/GO_MWU</u>.

## Author Contributions

BKL built and sequenced TagSeq libraries, and performed gene expression data analysis. WES performed the experiment which generated the samples. BKL and DIB wrote the manuscript. All authors approved the final version. The authors declare no conflicts of interest.

---

**Supplementary Table 1.**
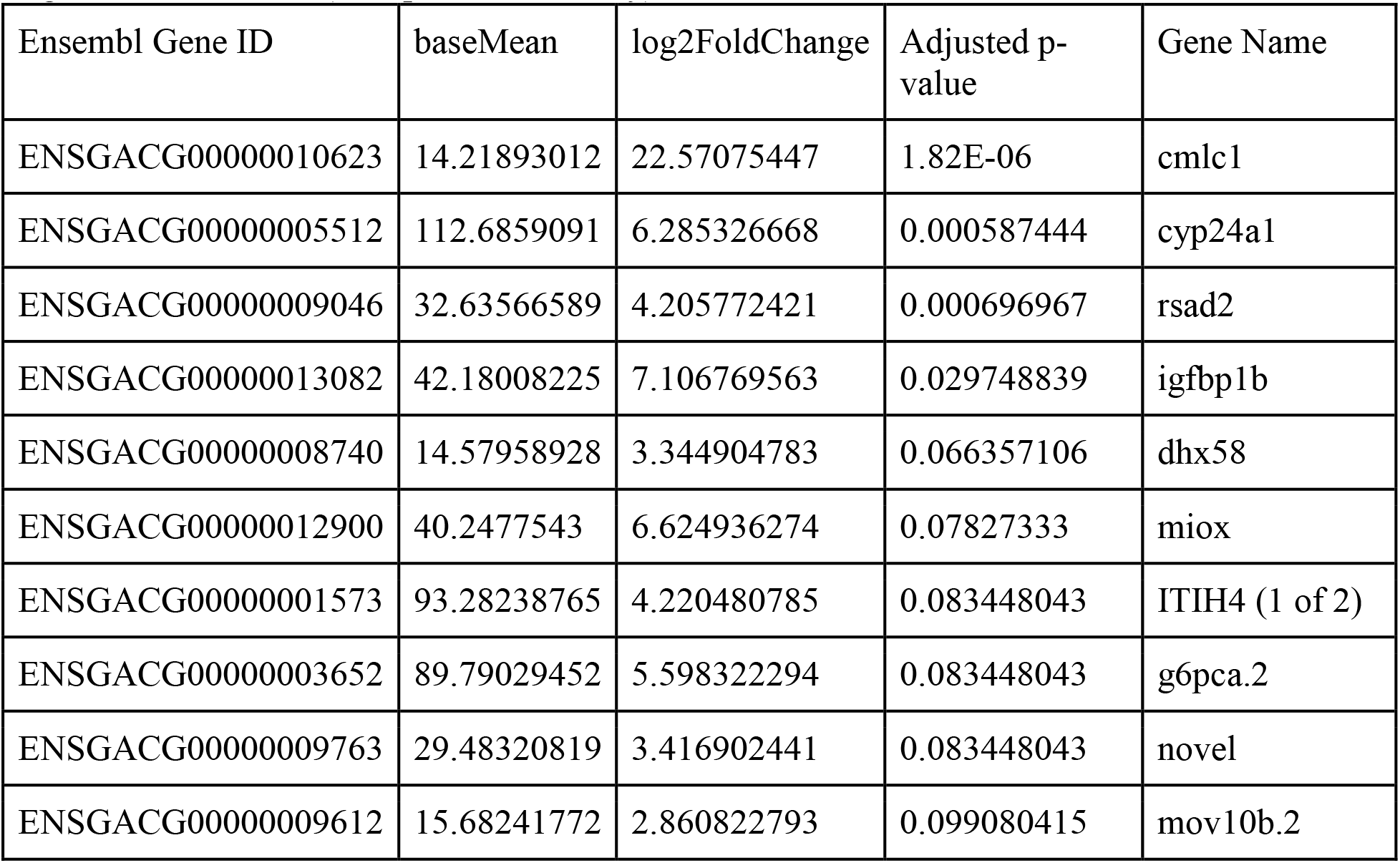
Summary statistics for genes significant for the interaction between origin and destination (transplanted fish only).

**Supplementary Figure 1.**
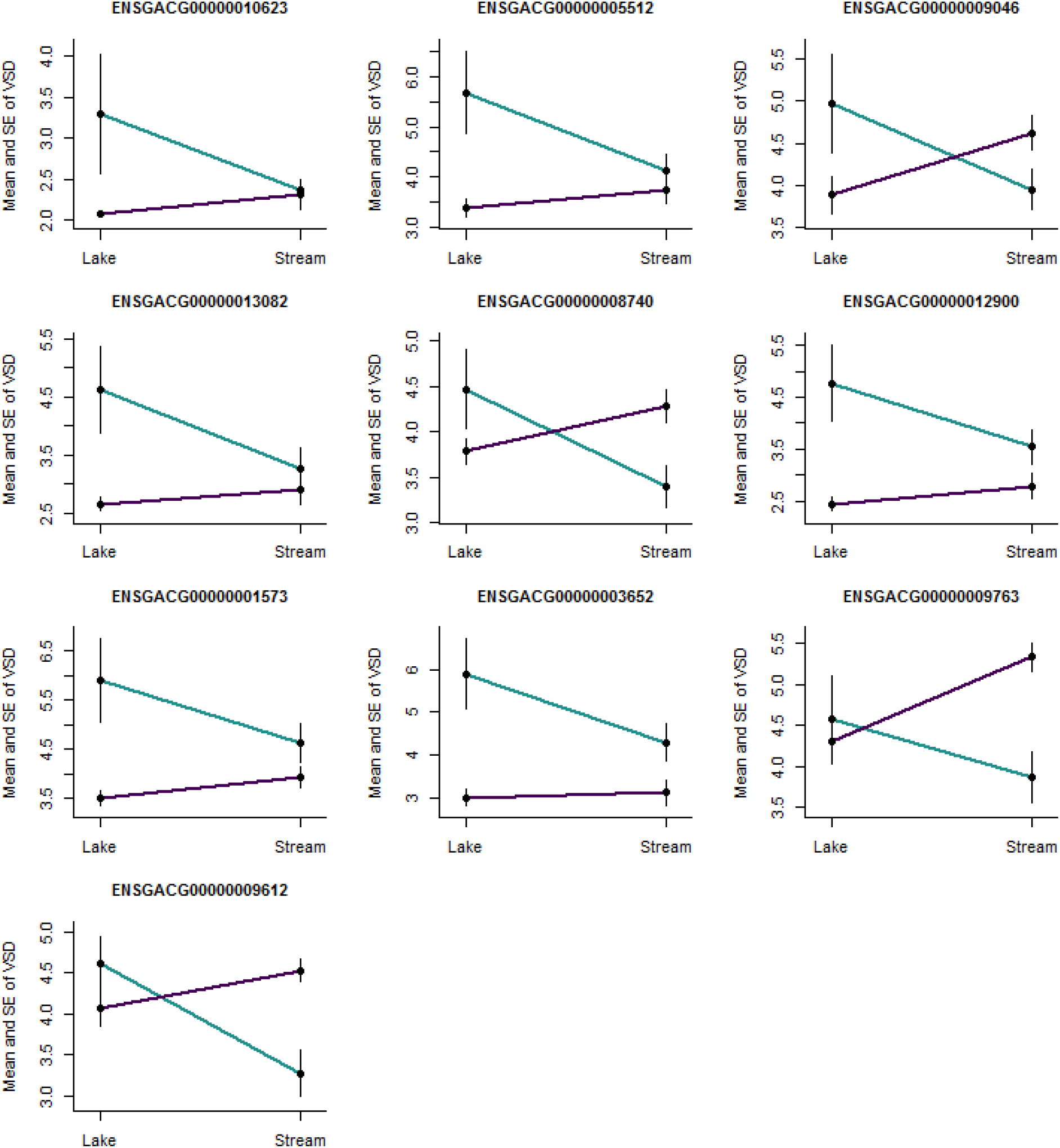
Reaction norms for all genes significant for the interaction between origin and destination. X axis is destination. Teal line indicates fish of lake origin and magenta line indicates fish of stream origin. Vertical lines indicate standard error.

**Supplementary Figure 2.**
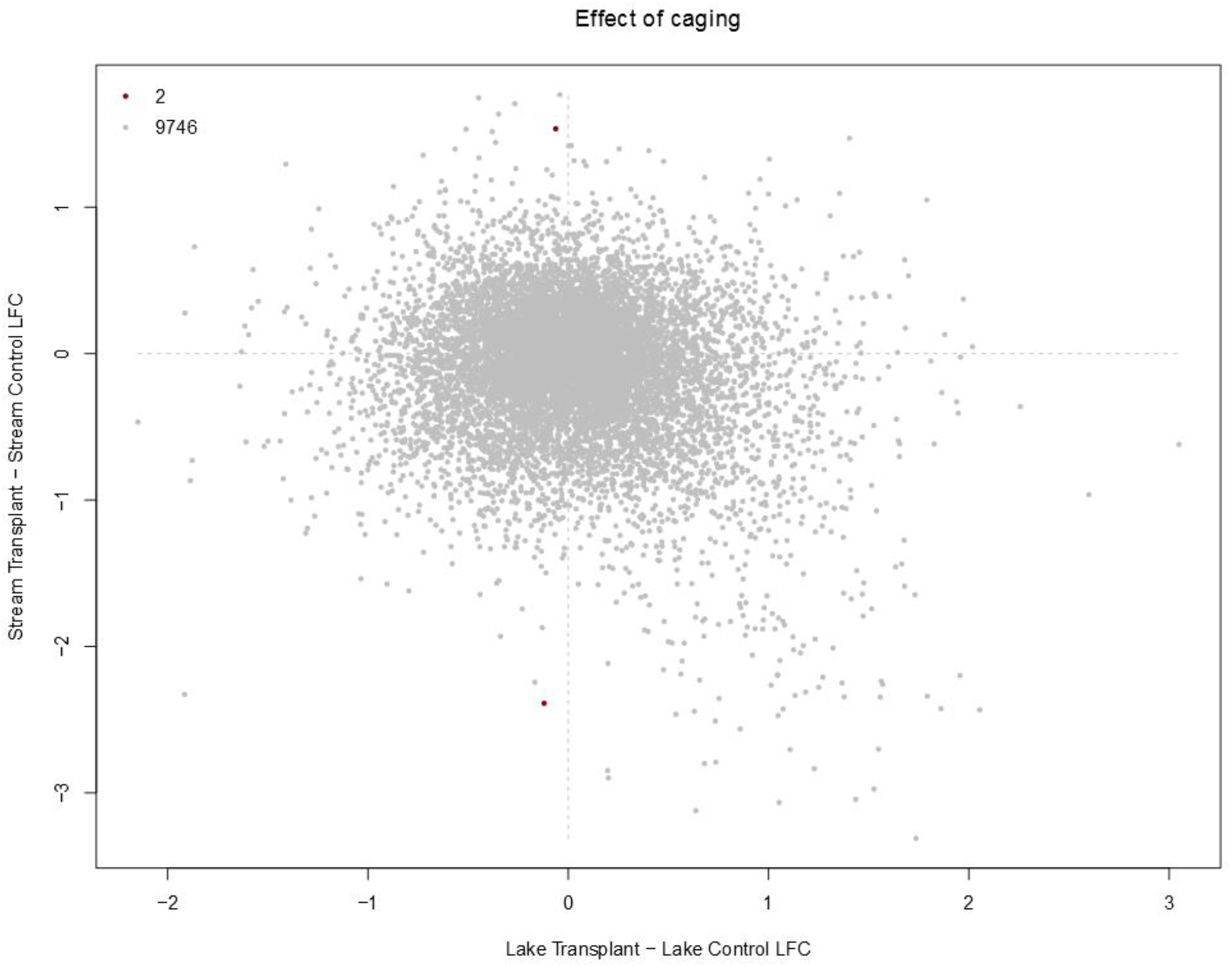
We sought to verify that our transplanted fish recapitulated the expression patterns of wild fish and that caging did not substantially alter our gene expression results on a gene by gene basis. Any generic response to caging should be observed in both the lake and in the stream. We included only wild fish and those that had been native transplants. We tested for a relationship between caging effect (LFC, native transplant - control) in lake and steam simultaneously with a linear model in limma (a single predictor with a level for each treatment) and contrasting control and natives within each habitat. We predicted that there should be no relationship between the cage effect in the lake versus the cage effect in the stream (i.e. this should produce a cloud of points about 0). Of the 9,748 gene we tested, only 2 differed significantly from this null expectation (q < 0.05).

**Supplementary Figure 3.**
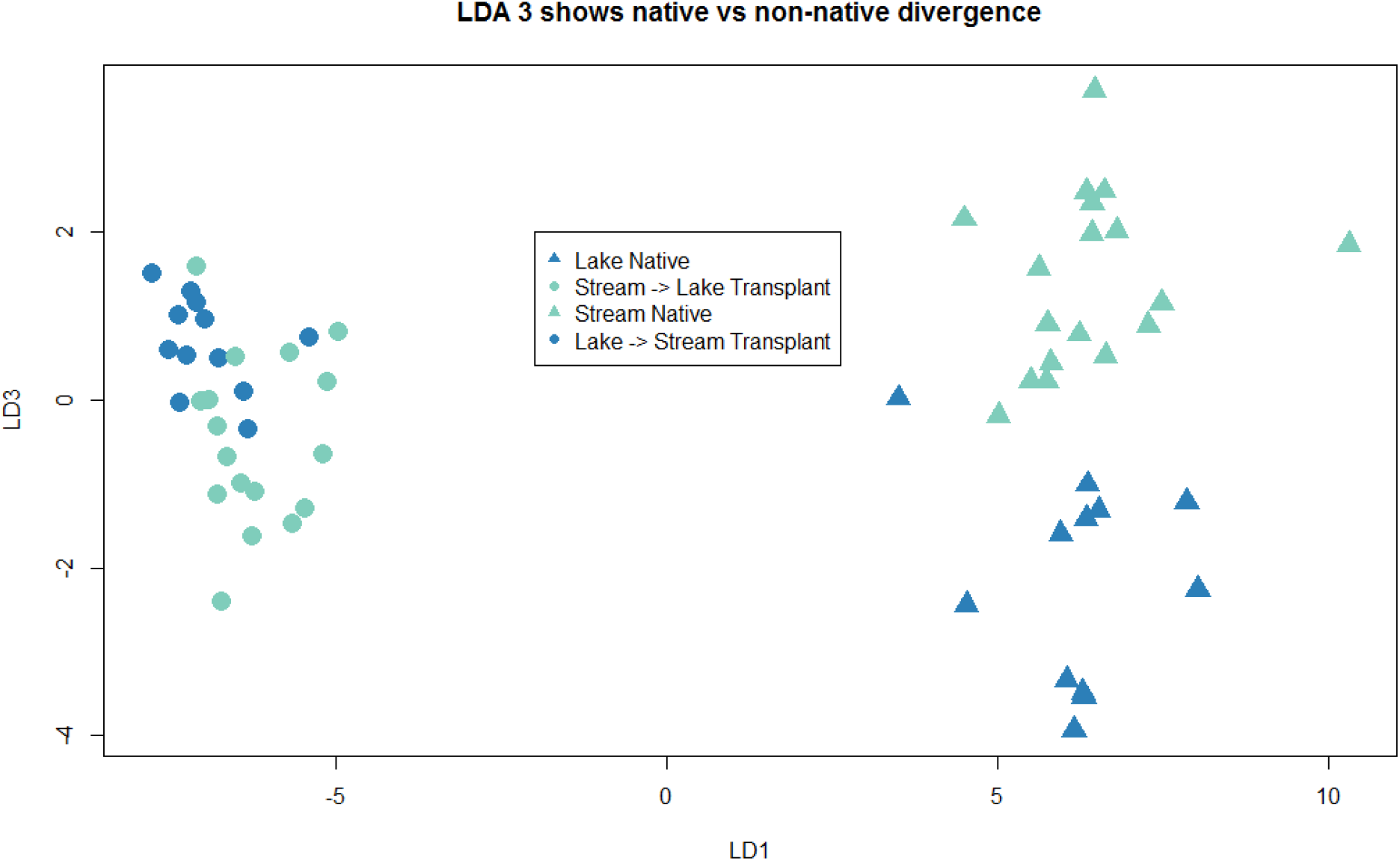
LD3 in convergence analysis roughly shows native vs immigrant differences in LD3.

